# Human Sound Localization Depends on Sound Intensity: Implications for Sensory Coding

**DOI:** 10.1101/378505

**Authors:** Antje Ihlefeld, Nima Alamatsaz, Robert M Shapley

## Abstract

A fundamental question of human perception is how we perceive target locations in space. Through our eyes and skin, the activation patterns of sensory organs provide rich spatial cues. However, for other sensory dimensions, including sound localization and visual depth perception, spatial locations must be computed by the brain. For instance, interaural time differences (ITDs) of the sounds reaching the ears allow listeners to localize sound in the horizontal plane. Our experiments tested two prevalent theories on how ITDs affect human sound localization: 1) the labelled-line model, encoding space through tuned representations of spatial location; versus 2) the hemispheric-difference model, representing space through spike-rate distances relative to a perceptual anchor. Unlike the labelled-line model, the hemispheric-difference model predicts that with decreasing intensity, sound localization should collapse toward midline reference, and this is what we observed behaviorally. These findings cast doubt on models of human sound localization that rely on a spatially tuned map. Moreover, analogous experimental results in vision indicate that perceived depth depends upon the contrast of the target. Based on our findings, we propose that the brain uses a canonical computation of location across sensory modalities: perceived location is encoded through population spike rate relative to baseline.

---

In the ascending mammalian auditory pathway, the first neural processing stage where ITDs are encoded, on the timescale of microseconds, is the medial superior olive (MSO). Here, temporally precise binaural inputs converge, and their ITDs are converted to neural firing rate (Goldberg and Brown 1969; Yin and Chan 1990; Spitzer and Semple 1995; Pecka et al., 2010; Day and Semple 2011). The shape of the MSO output firing rate curves as a function of ITD resembles that of a cross-correlation operation on the inputs to each ear (Batra and Yin 2004). How this information is interpreted downstream of the MSO has led to the development of conflicting theories on the neural mechanisms of sound localization in humans. One prominent neural model for sound localization, originally proposed by Jeffress, consists of a labelled line of coincidence detector neurons that are sensitive to the binaural synchronicity of neural inputs from each ear (Jeffress, 1948), with each neuron maximally sensitive to a specific magnitude of ITD (Figure 1A). This labelled-line model is computationally equivalent to a neural place code based on bandlimited cross-correlations of the sounds reaching both ears (Domnitz and Colburn, 1977). Several studies support the existence of labelled-line neural place code mechanisms in the avian brain (Carr and Konishi, 1988; Overholt et al., 1992), and versions of it have successfully been applied in many engineering applications predicting human localization performance (e.g., Durlach, 1963; Hafter, 1971; Stern and Trahiotis, 1995; Breebaart et al., 2001; Hartmann et al., 2005).

**Fig 1.**
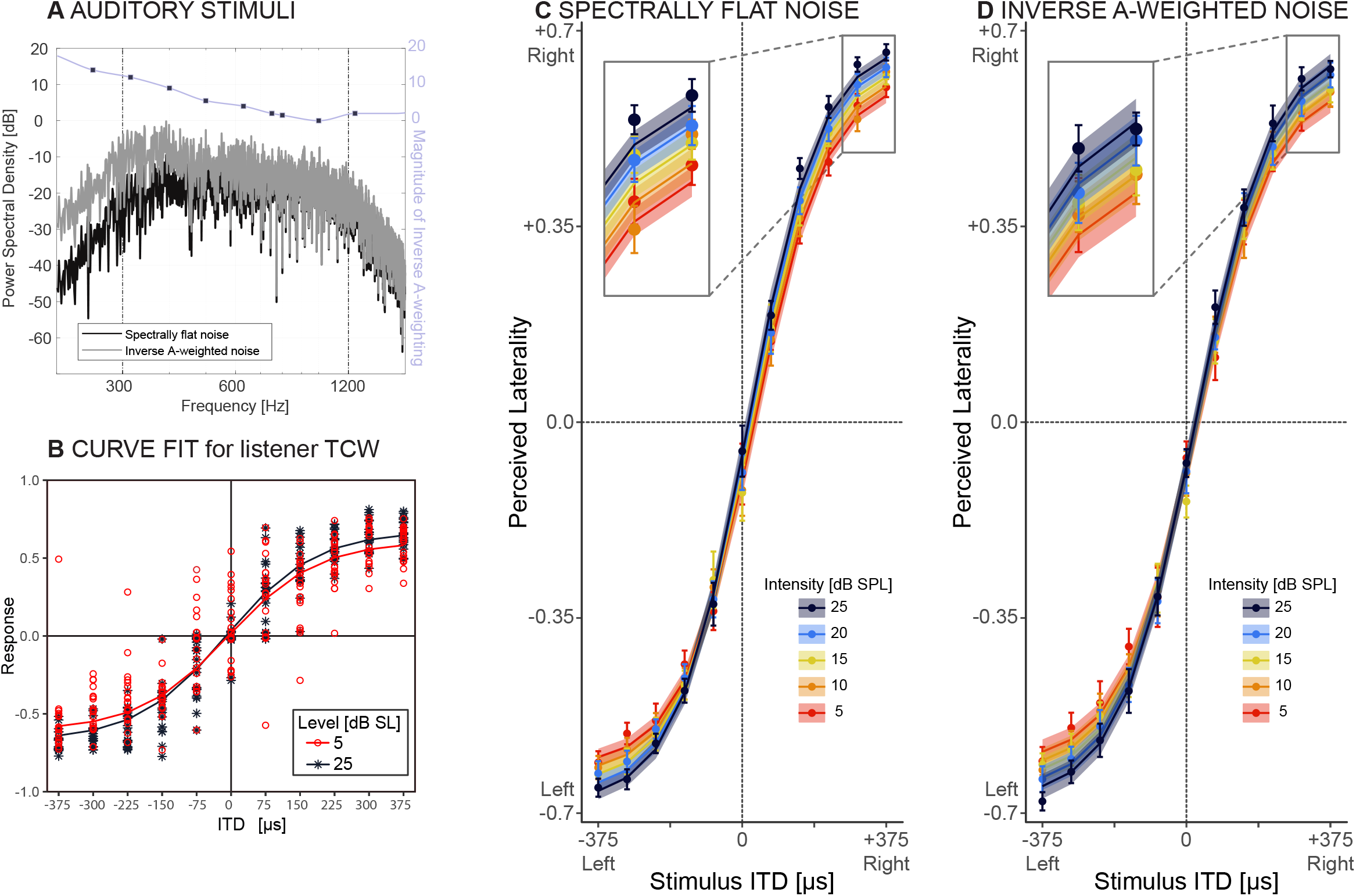
A) Firing rate of a simulated *nucleus laminaris* neuron with a preferred ITD of 375 μs, as a function of source ITD. The model predicts source laterality based on the locus of the peak of the firing rate function. B) Hemispheric differences in firing rates, averaged across all 81 simulated *inferior colliculus* units. Rate models assume that source laterality is proportional to firing rate, causing ambiguities at the lowest sound intensities. Inset: Reconstructed responses of an *inferior colliculus* unit. The unit predominantly responds contralaterally to the direction of sound (high-contrast traces). The hemispheric difference model subtracts this activity from the average rate on the ipsilateral side (example shown with low-contrast traces). C) Mean population response using labelled-line coding across a range of ITDs and sound intensities. Inset: The root-mean square (RMS) difference relative to estimated angle at 80 dB SPL does not change with sound intensity, predicting that sound laterality is intensity invariant. D) Mean population response using hemispheric-difference coding. For lower sound intensities, predicted source direction is biased towards midline (compare red and orange versus blue or yellow). For higher sound intensities, predicted source direction is intensity invariant (blue on top of yellow line). Inset: RMS difference relative to estimated angle at 80 dB SPL decreases with increasing sound intensity, predicting that sound laterality is not intensity invariant. Ribbons show one standard error of the mean across 100 simulated responses. Sound intensity is denoted by color (see color key in the figure).

A growing literature proposes an alternative to the labelled-line model to explain mammalian sensitivity to ITD. One reason for an alternative is that two excitatory inputs should suffice to implement the labelled-line model, but evidence from experiments on *Mongolian gerbils* shows that in addition to bilateral excitatory inputs, sharply tuned bilateral inhibitory inputs to the MSO play a crucial role in processing ITDs (Brand, et al., 2002). Moreover, to date no labelled-line type neurons have been discovered in a mammalian species. Indeed, using a population rate code, several studies proposed that mammalian sound localization can be modeled based on differences in firing rates across the two hemispheres (Figure 1B; van Bergeijk. 1962; McAlpine and Grothe, 2003; Devore et al., 2009). Rate-based models generally predict that neuronal responses carry most information at the steepest slopes of neural-discharge-rate versus ITD curves, where neural discharge changes most strongly (Stecker et al., 2005), consistent with the observation that the peak ITDs of rate-ITD curves often fall outside the physiologically plausible range (McAlpine and Grothe, 2003; Grothe et al., 2010; but see also Joris et al., 2006). In addition, some authors have suggested that the findings that mammalian sound localization can adapt to stimulus history are further support for a rate-based neural population code (Phillips and Hall, 2005; Stange et al., 2013).

It is unknown which of the two competing models, broadly characterized as labelled-line versus rate-code model, describes human sound localization better. Here, we observe that the two different models predict different dependencies of sound localization on sound intensity. By combining behavioral data on sound intensity dependence in normal-hearing listeners with numerical predictions of human sound lateralization from both models, we attempt to disentangle whether human auditory perception is based on a place code, akin to the labelled-line model, or whether it is instead more closely described by a population rate code.

To predict how lateralization depends on sound intensity from the responses of labelled-line neurons, we estimated neural firing rates from previous recordings in the *nucleus laminaris* in *barn owl* (Peña et al., 1999). To estimate lateralization’s dependence on level based on a population rate code, we used previous recordings from the *inferior colliculus* of *rhesus macaque monkey* and calculated hemispheric differences in firing rate (Zwiers et al., 2004). The labelled-line neurons predicted that, as sound intensity decreases, perceived source laterality would converge towards similar means for low versus high sound intensities, with increased response variability at decreasing sound intensities (Figure 1C). In contrast, the hemispheric-difference model predicted that as sound intensity decreases to near threshold levels, perceived laterality would become increasingly biased toward the midline reference (Figure 1D). At higher overall sound intensities, both models predicted that lateralization would be intensity invariant (see insets in Figure 1C versus D). Therefore, analyzing how sound intensity affects perceived sound direction near sensation threshold offers an opportunity to disentangle whether our human auditory system relies on a place-based or rate-based population code for localizing sound based on ITD.

A listener’s ability to discriminate ITD can vary with sound intensity (Dietz et al., 2013). However, it is difficult to interpret previous findings linking ITD and sound localization as a function of sound intensity. Some reported decreased lateralization near sensation threshold (Teas, 1962, Sabin et al. 2005), but others reported weak or no level effects on lateralization (von Békésy and Wever, 1960; Mickunas 1963; Hartmann and Rakerd 1993; Macpherson and Middlebrooks 2000; Inoue 2001; Vliegen and Van Opstal 2004; Brungart and Simpson 2008; Gai et al., 2013). Several factors complicate the interpretation of these previous findings in the context of the current hypothesis. For instance, assuming an approximately 30 dB dynamic range of rate-level function either at the MSO or downstream in the binaural pathway (e.g., medial superior olive: Goldberg and Brown, 1969; inferior colliculus: Zwiers et al., 2004), for stimuli at higher sensation levels (SL) where the rate-level functions saturate, both the labelled-line and the hemispheric difference model predict level invariance. This could explain how studies that tested for sound intensity effects over a range of high intensities did not see an effect. Moreover, when presented in the free field, in addition to ITD, sounds also contain interaural level differences (ILDs) and spectral cues. For low-frequency sound, listeners rely dominantly on ITD when judging lateral source angle (Strutt, 1907). However, for broadband sound, listeners integrate across all three types of spatial cue (Wightman and, 1992; Ihlefeld and Shinn-Cunningham, 2011). Unlike ITDs, ILDs and overall sound intensity both decrease with increasing source distance, raising the possibility that for stimuli with high-frequency content, listeners judged softer sounds to be more medial because they interpreted them to be farther away than louder sounds. Further, at low sound intensities, the sound-direction-related notches of the spectral cues at high-frequencies should have been less audible than at higher sound intensities, increasing stimulus ambiguity. A resulting increase in response variability may have obscured the effect of level on ITD coding. Finally, some historic studies used only two or three listeners, suggesting that they may have been statistically underpowered. Thus, the literature provides insufficient evidence on how ITD-based lateralization varies with level near sensation threshold.

Here, we tested the null hypothesis that ITD-based human sound localization relies on a population rate-place neural code. This hypothesis predicted that the mean perceived direction based on ITD would be intensity invariant, even at intensities close to SL. Using a psychophysical paradigm, we studied lateralization based on ITD as a function of sound intensity in a group of ten normally hearing listeners (experiment 1). Stimuli consisted of low-frequency noise tokens that were bandlimited to cover most of the frequency range where humans can discriminate ITD (Brughera et al 2013; here, corner frequencies from 300 to 1200 Hz, shown in Figure 2A). In each one-interval trial, listeners had to indicate perceived laterality across a range of ITDs from -375 to 375 μs. Lateralization was measured as function of SL. To examine how sound intensity affects perceived ITD coding of source direction, we modelled perceived laterality with a nonlinear mixed effect model (NLME) that included fixed effects of ITD and sound intensity as well as a random effect of listener.

**Fig 2.**
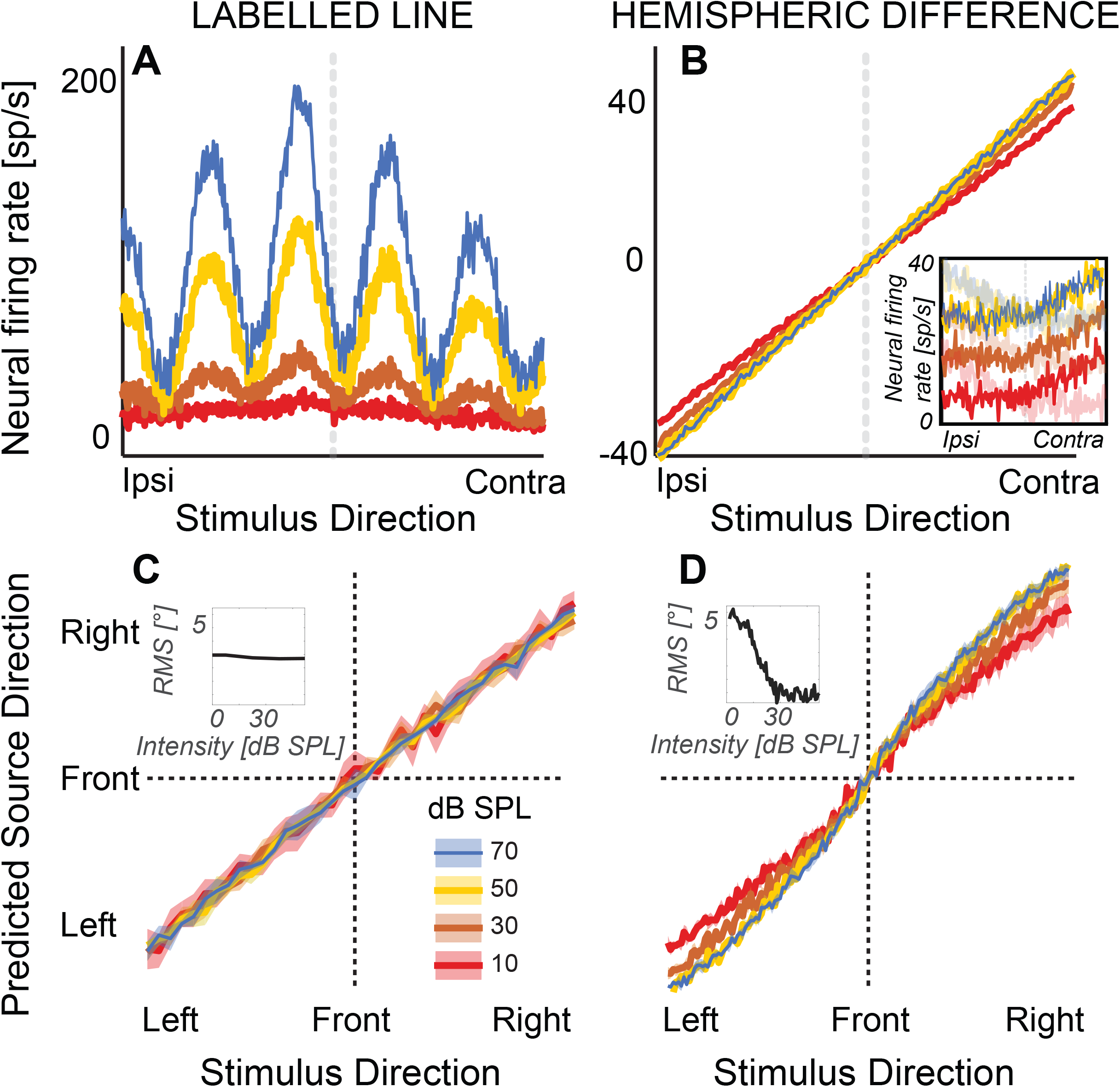
Behavioral experiment A) Stimuli: spectrally flat noise, used in experiment 1 (dark grey) versus A-weighted noise, tested as a control for audibility in experiment 2 (light grey). The purple line shows the magnitude of the zero-phase inverse A-weighting filter. B) Responses from one representative listener (TCW) across two sound intensities and the corresponding NLME fits for these data. C and D) Perceived laterality as a function of ITD for C) spectrally flat noise (experiment 1) or D) A-weighted noise (experiment 2). Error bars, where large enough to be visible, show one standard error of the mean across listeners. Colors denote sound intensity. Insets illustrate magnified section of the plots. Circles show raw data, lines and ribbons show NLME fits and one standard of the mean.

Figure 2B depicts lateralization performance with spectrally flat noise at two sound intensities for a representative listener (TCW). Figure 2C shows raw data (circles) and NMLE fits (lines) across all listeners. Error bars show one standard error of the mean across listeners, and shaded ribbons indicate one standard error of the mean fit across listeners. This model predicts 82.7% of the variance in the measured responses and is deemed an appropriate fit of the data. Table I lists all NLME parameters. Perceived laterality scores increased with increasing ITD, as expected.

**Table I.** Results of Non-linear Mixed Effects Model for flat-spectrum noise condition.

|  | Value | Std.Error | df | t.value | p.value |
| --- | --- | --- | --- | --- | --- |
| $\alpha_{x0}$ [1/μs] | | | | | |
| *Slope attributed to ITD* | 2.45 | 0.05 | 10988 | 46.10 | <0.001 |
| $\alpha_{x1}$ [1/dB] | | | | | |
| *Slope attributed to sound intensity* | 0.02 | 0.01 | 10988 | 1.46 | 0.14 |
| $\alpha_{y1}$ [1/dB] | | | | | |
| *Lateralization attributed to sound intensity* | 0.05 | 0.01 | 10988 | 7.58 | <0.001 |

With decreasing sound intensity, percepts were increasingly biased towards midline (compare order of colored lines, magnified in the inset of Figure 2C). These trends were supported by the NLME model, which revealed significant effects of ITD (*α*_*x*0_ *estimate* = 2.45, *SE* = 0.05, *p* < 0.001) and sound intensity (*α*_*y*1_ *estimate* = 0.05, SE = 0.01, p < 0.001), rejecting our null hypothesis. Varying the coefficient modelling the slope of perceived laterality as a function of level did not alter the model fit significantly (*α*_*x*1_ estimate = 0.02, SE = 0.01, p = 0.14).

In a second experiment, we examined whether these results were robust to the spectral details of the stimuli. A caveat of testing spectrally flat noise at low sound intensities is that parts of the spectrum may be inaudible. Therefore, the results of experiment 1 could potentially be confounded by the fact that the bandwidth of the audible portion of the noise tokens decreased with decreasing sound intensity.

Therefore, as a control for stimulus audibility, the same listeners were tested again, using inverse A-weighted noises (experiment 2). All of the original ten listeners from experiment 1 completed experiment 2. Methods were similar as in the first experiment, except that the stimuli consisted of inversely A-weighted noise (compare magnitude spectra in Figure 2A). The data and NLME model fits for the second experiment are shown in Figure 2D (color key identical to Figure 2C), and coefficients are listed in Table II. This second model accounts for 82.8% of the variance in the data, closely fitting the measured responses. All NLME coefficients are significant (*α*_*x*0_ *estimate* = 2.57, *SE* = 0.06; *α*_*x*1_ *estimate* = 0.07, *SE* = 0.01; *α*_*y*1_ *estimate* = 0.04, *SE* = 0.01, *p* < 0.001). The fact that *α*_*x*1_ is significant shows that when all noise portions are approximately equally audible, as here, with inverse A-weighted noise, both perceived laterality and the slope linking the change in laterality to ITD decrease with decreasing sound intensity. Thus, the results confirm the effect of biasing perceived laterality toward midline with decreasing sound intensity. Therefore, for both spectrally flat noise and A-weighted noise, statistical analyses, which partialed out overall differences between listeners, are inconsistent with a labelled-line model of human sound localization. The results challenge the view that human sound localization is encoded through a rate-place population code.

**Table II.** Results of Non-linear Mixed Effects Model for inverse A-weighted noise condition.

|  | Value | Std.Error | df | t.value | p.value |
| --- | --- | --- | --- | --- | --- |
| $\alpha_{x0}$ [1/μs] | | | | | |
| *Slope attributed to ITD* | 2.57 | 0.06 | 10988 | 46.24 | <0.001 |
| $\alpha_{x1}$ [1/dB] | | | | | |
| *Slope attributed to sound intensity* | 0.07 | 0.01 | 10988 | 5.00 | <0.001 |
| $\alpha_{y1}$ [1/dB] | | | | | |
| *Lateralization attributed to sound intensity* | 0.04 | 0.01 | 10988 | 7.0 | <0.001 |

Population rate coding to compute sensory dimension may not be unique to the auditory system. In analogy to sound localization based on the comparison of signals from the two ears (Figure 3A), visual depth is computed in the cerebral cortex based on signals from the two eyes (Figure 3B; Poggio 1995; Parker and Cumming 2001; Parker 2016). Specifically, in both primary *V1* and extrastriate *V3a* cortex of *rhesus macaque* monkeys, three types of neurons are thought to encode binocular disparity. “Tuned-excitatory” neurons respond best to zero spatial disparity between the two eyes, whereas “near cells” responds more vigorously when an object approaches, increasing crossed disparity between the eyes (Parker and Cumming 2001). Finally, “far cells” fire more vigorously as uncrossed disparity increases. In *V1*, the most frequently encountered type of binocular neurons are of the tuned-excitatory type. However, in *V3a* the large majority of neurons is stereo-specific (Poggio et al 1988) and most neurons are either near or far cells. Functional magnetic resonance imaging experiments on human stereoscopic vision found that unlike *V1* activity, the activity in cortical area *V3a* predicts behavioral performance on tasks involving stereoscopic depth (Backus et al., 2001). Thus, we propose that near and far cells encode visual distance from the fixation plane in a way similar to how *inferior colliculus* neurons encode auditory azimuthal angle away from midline reference: firing rate increases monotonically with distance from perceptual reference anchor or fixation.

**Fig 3.**
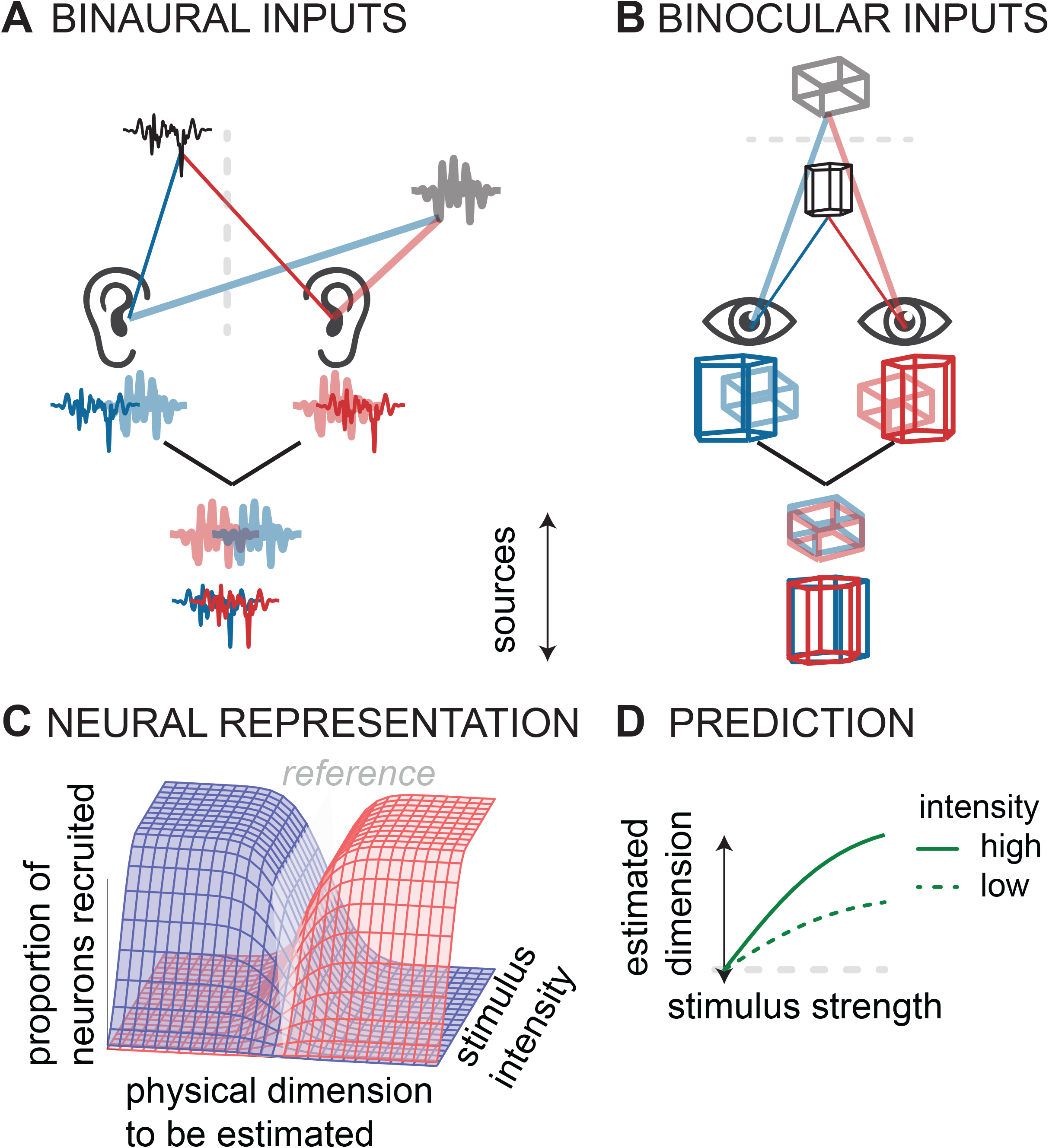
A) Computing sound direction requires analysis of the binaural difference between the signals reaching the left and right ear. B) Estimating visual depth hinges on analysis of the binocular disparity between the signals reaching left and right eye. C) For both hearing and vision, the proportion of the neural population that is stimulated (in the *inferior colliculus* or *V3)* depends both on the physical dimension to be estimated (source laterality or source distance) and the intensity of the stimulus (sound intensity or visual contrast). For hearing and vision, ambiguity in this putative neural code predicts D) biased responses at low stimulus intensities (sound intensity or contrast).

We observe that in both the auditory and the visual system, the same cells that are tuned to binaural ITD or binocular disparity also have intensity-response functions. A rate code based on a population of these cells should cause ambiguities when stimulated below the saturation firing rate, either at low sound intensity or at low contrast (Figure 3C). Thus, based on the analogies between the stereo-depth computation and the azimuth-ITD computation, we hypothesized that low visual contrast might affect the computation of depth in a manner analogous to the effect of low sound levels in sound localization—there might be a bias to lower perceived depth at lower contrast (Figure 3D). Indeed, one study found such an effect, but only in some observers (Cisarik and Harwerth 2008). A confounding factor in that earlier study is that perceived depth is a complicated neural computation, not only dependent on stereoscopic disparity but also on monocular cues including contrast (Parker 2016). Several studies on depth perception indicate that low contrast is interpreted by the brain as a cue for distance; lower contrast targets are perceived farther away (e.g. Schor and Howarth, 1986; Rohaly and Wilson, 1999). However, elegantly designed experiments that were not affected by the low contrast bias demonstrated that low contrast causes perceived depth to shrink, both for near and far deviations from baseline (Chen et al., 2016). Thus, there is a link between population rate coding and stimulus intensity in perceived visual depth as in perceived auditory azimuth, two perceptual spatial dimensions computed by the brain.

In summary, unlike predictions from a hemispheric difference model, labelled-line coding predicts that sound localization is intensity invariant. Our experimental results show that for low frequency noise, where ITDs are the dominant localization cue, and at low sound intensities, sound lateralization based on ITD is not intensity invariant; it becomes increasingly medially biased with decreasing SL. This finding parallels a phenomenon of fixation bias when calculating visual distance from binocular disparity at low contrast. This casts doubt on the idea that the neural mechanism of ITD-based sound localization and binocular disparity-based visual distance estimation are based on place-based coding. Instead, our perceptual data on auditory localization together with previously published data on visual distance perception are parsimonious with the idea that a population rate code underlies the brain’s computation of location.

## METHODS

### 1. Listeners

Twelve naïve normal-hearing listeners (ages 18-27, five females) were enrolled in this study and paid for their time. Their audiometric thresholds, as assessed via a calibrated GSI 39 Auto Tymp device (Grason-Stadler), were 25 dB hearing level or better at octave frequencies from 250 to 8000 Hz, and did not differ by more than 10 dB across ears at each octave frequency. All testing was administered according to the guidelines of the Institutional Review Board of the New Jersey Institute of Technology.

### 2. Overall design

Listeners were seated in a double-walled sound-attenuating booth (Industrial Acoustics Chamber) with a noise floor of 20.0 dB SPL (wideband L_AFeq_). Stimuli were digitally generated in Matlab R2016b (The MathWorks, Inc.), D/A converted through an external sound card (Emotiva Stealth DC-1) at a sampling frequency of 192 kHz, with a resolution of 24 bits per sample, and presented to the listener through ER-2 insert earphones (Etymotic Research Inc.). The equipment was calibrated using an acoustic mannequin (KEMAR model, G.R.A.S. Sound and Vibration) with a precision of less than 10 μs ITD and less than 2 dB ILD. Foam eartips were inserted following guidelines provided by Etymotic Research to encourage equal representation of sounds to both ears (no ILD) and minimize interaural leakage. Each session lasted approximately 60 minutes. Listeners kept the insert earphones placed inside their ears throughout testing. Insert earphones were replaced by the experimenter after each break.

Throughout this study, to generate stimuli, tokens of uniformly distributed white noise were generated and bandpassed using a zero-phase Butterworth filter with 36 dB/octave frequency roll-off, and 3 dB down points at 300 and 1200 Hz. Each noise token was 1 s in duration, including 10 ms long squared cosine ramps at the onset and offset.

### 3. Sensation level measurements

At the beginning of each session, and, as a re-test control, mid-way through each session, each individual listener’s SL was measured for the type of sound that was later on used for training and testing, via one run of adaptive tracking. On each one-interval trial of each track, a new noise token was generated and presented diotically. Trials were spaced randomly in time (uniform distribution, inter-token intervals from 3 to 5.5 s). Listeners pressed a button when they heard a sound. No response feedback was given.

On each trial, a response was scored a “hit” if a listener responded with a button push before the onset of the subsequent trial, and a “miss” if the listener did not respond during the interval. If a listener’s response changed from hit to miss or from miss to hit across sequential trials, this was interpreted as a response reversal. Using one-up-one-down adaptive tracking, the noise intensity was increased or decreased after each reversal, with a step size of 5 dB (decreasing) or 2.5 dB (increasing). Each listener completed ten adaptive-track reversals, with SL threshold equaling the median of the final six reversals. Each SL was used as reference intensity for the subsequent 30 minutes of testing. If detection thresholds changed between initial test and re-test control by more than 5 dB, this indicated that an insert earphone moved, and the experimenter replaced the earphones. Thresholds generally did not change by more than 5 dB.

### 4. Training

To train listeners on consistently reporting their perception of ITD, using adaptive tracking, listeners matched the perceived laterality of a variable-ITD pointer to that of a fixed-ITD target. Target token intensity was set relative to the listener’s own diotic sensation threshold, at 10 or 25 dB SL, and presented with 0 dB ILD. The pointer intensity was fixed at 25 dB SL. Target ITDs spanned the range from -375 to 375 μs, in 75 μs steps. Target ITDs and SLs were randomly interleaved across runs, but held fixed throughout each adaptive run.

In each two-interval trial of a run, the pointer token was presented in the first, and the target token in the second interval. The start ITD of the pointer token at the beginning of each run equaled 0 μs. Using a hand-held controller (Xbox 360 wireless controller for Windows, Microsoft Corp.), listeners adjusted the ITD of the pointer token. Specifically, listeners pushed the directional keys (D-pad) either to the left or right in order to move and match the pointer direction with that of the target sound. When a listener indicated a left- or right-ward response, the pointer ITD was decreased or increased. Initial ITD step size equaled 100 μs, then 50 ± 5 μs (uniformly distributed) after the first reversal. By the end of the second reversal, ITD step size was reduced to 25 ± 5 μs (uniformly distributed) and remained the same for all of the following reversals. Listeners were instructed to “home in” on the target by moving the pointer initially to a position more lateral than the target, then more medial than the target with the goal of centering on the target. No response feedback was provided. A run was completed after a listener had completed a total of five adaptive-tracking reversals. For each target ITD, the matched pointer ITD was estimated by averaging the pointer ITDs of the final two reversals.

Each listener performed three sessions of training: In the first session only a subset of target ITDs were presented (-375, -150, 0, 150 and 375 μs), whereas the two following sessions included all of the eleven ITDs. Per training session, each ITD was presented once at 10 and 25 dB SL, for a total of 54 adaptive tracking runs across all training sessions. To familiarize listeners with the experimental task (described below), at the end of second and third sessions of training listeners performed an additional 5 blocks of the experimental testing task, without response feedback. These task training data were not used for statistical analysis.

To assess whether listeners could reliably report their lateralization percepts, training performance was evaluated for each listener by calculating the Pearson correlation coefficient between target ITD and matched pointer ITD in the final training session. Criterion correlation equaled 0.9 (N=11 ITDs, significance level=0.01, power=0.95). Ten listeners reached criterion, suggesting that they were able to consistently report where they perceived the sounds based on ITD. Two of the originally recruited twelve listeners failed to reach training criterion (R^2^=0.84, 0.87) and were excluded from testing.

### 5. Testing

Using the method of fixed stimuli, we tested lateralization in two experiments. Except for the stimuli, which consisted of spectrally flat noise tokens in experiment 1 and A-weighted noise tokens in experiment 2, the methods were similar across the two experiments. Noise tokens were generated from a statistically similar noise distribution as those presented during both SL measurements and training (see Overall Design). A touchscreen monitor (Dell P2314T) displayed the response interface at about 40 cm distance from the listener. Using a precise touch stylus (MEKO Active Fine Point Stylus 1.5 mm Tip), listeners indicated perceived laterality of noise in a one-interval task. Noise tokens were presented at 5, 10, 15, 20, and 25 dB SL. ITDs varied randomly from trial to trial, in 75 μs steps spanning the range from -375 μs to +375 μs. On each trial, a new token of noise was generated. Each listener performed 20 blocks of 55 trials each (11 ITDs at each of the 5 sound intensities), with SL measured both before the first and the eleventh block. ITDs and sound intensity were randomly interleaved from trial to trial such that each combination of ITD and sound intensity was presented once before all of them were repeated in a different random order.

### 6. Statistical Analysis

Growth curve analysis was used to analyze perceived laterality scores as a function of ITD and sound intensity. For each of the two noise conditions, the perceived laterality scores were fitted with an NLME model. The model included fixed effects *α* and random effects *β*. Equation 1 describes a sigmoidal function linking ITD to perceived laterality, with a score from left (-1) to right (1). The effect of sound intensity on the maximal extent of lateralization is *α*_*y*1_, the slope terms are *α*_*x*0_ for perceived laterality changes attributed to ITD, and *α*_*x*1_ for laterality-ITD slopes attributed to sound intensity. Random effects of individual differences across listeners were used to model both the maximal extent of lateralization, *β*_*y*0,*listener*_, and the perceived midline, *β*_*x*0,*listener*_, centering the sigmoid.

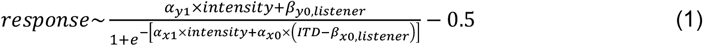

To better conform with the assumptions of the NLME model, prior to fitting, ITD and sound intensity parameters were scaled by subtracting the mean stimulus value, and dividing by the maximal stimulus value, resulting in distributions of stimulus parameters with zero-mean and a variance of one. Laterality scores were then fitted using these normalized parameters, with the nlme package, programmed in RStudio 1.1 for Windows (RStudio Inc., Boston, MA, USA).

### 7. Models of neural coding

We estimated the combined effects of ITD and sound intensity on predicted source laterality both in avian labelled-line type units and in binaurally sensitive units of a mammalian auditory system. Few studies have measured neural discharge rate as a function of ITD at very low sound intensities. However, one study in barn owl shows that the output functions of *nucleus laminaris* neurons can be modeled through interaural cross-correlation functions, even at very low sound intensities (Peña et al., 1996). To mimic the type of auditory information available at the output of a cross-correlator neuron, for each source ITD, we added dichotic noise tokens (0 dB ILD) and amplitude weighted dichotic noise. We then processed this partially interaurally correlated mixture with 1/3-octave wide bandpass filters of 1 kHz center frequency and 24 dB/octave frequency roll-off. We scaled the dichotic noise tokens such that the resulting Pearson correlation coefficients of the interaural cross-correlation functions matched those previously reported in the *nucleus laminaris* of barn owl with a precision error of less than 10% (Peña et al., 1996). This labelled-line model then predicted source laterality through the ITD corresponding to the peak of the resulting cross-correlation function. To estimate the mean and variance of predicted ITD as a function of sound intensity, we ran 100 simulations.

In the mammalian auditory system, one previous study reports firing statistics for 81 *inferior colliculus* units in *rhesus macaque* as a function of ITD and over a wide range of sound intensities, including very low sound intensities (Zwiers et al., 1999). To estimate firing rate as a function of ITD and sound intensity in the *inferior colliculus*, we used previously published regression parameters linking ITD, sound intensity and firing rate through linear regression fits and assumed that each *inferior colliculus* unit fired with a sigmoidal output function that saturated over a 30 dB dynamic range, had linear growth over the physiologically plausible range of ITDs, mostly responded contra-laterally, had a threshold between 0 and 10 dB SPL, and a spontaneous non-sound-evoked discharge of 5 spikes/second (e.g., Ramachandran et al., 1999). We then used these firing rates to estimate, collapsed across sound intensities from 0 to 80 dB SPL, the probability density of the firing rate for each *inferior colliculus* unit as a function of source ITD, and estimated the sound azimuth as a function of sound intensity via maximum likelihood. To estimate the mean and variance of predicted ITD as a function of sound intensity, we then ran a bootstrapping analysis, sampling with replacement 100 times.

## ACKNOWLEDGMENTS

The authors thank Joshua Hajicek for sharing code for sensation threshold measurements, and Jaasrini Vellore and Patrycja Puzio for helping with data collection. This work was funded by a start-up grant to AI at NJIT.

